# Psychoacoustic Study of the Rock Art Sites of Cuevas de la Araña (Bicorp, Spain)

**DOI:** 10.1101/2025.01.14.629613

**Authors:** Samantha López, Tapio Lokki, Margarita Díaz-Andreu, Carles Escera

## Abstract

Acoustics play a crucial role in shaping our perception of sound and its emotional impact. The rock art site of *Cuevas de la Araña* in Bicorp, Spain, is an archaeological site where pre-historic communities gathered for social and ritual activities. *Cuevas de la Araña* exhibits acoustical characteristics that could have enhanced the sensory and emotional impact during ceremonies performed with music. In the present study, a listening test was conducted to assess how the acoustics of *Cuevas de la Araña* and other rock art sites influence modern-day listeners’ perception of sound. Listeners were asked to describe, using their own vocabulary, a series of auralizations created with the impulse responses collected in *Cuevas de la Araña* and other neighboring sites with and without rock art. The words written by participants underwent categorization through a hierarchical clustering approach. Significant results emerged indicating that listeners perceived auralizations from rock art sites as larger, wider, less direct, farther and more reverberant than the auralizations from sites lacking rock art. Notably, the most prominent disparities were observed in the categories of size, distance, and reverberation when contrasting the auralizations from *Cuevas de la Araña* with those of non-painted sites. These findings align with the outcomes documented in prior literature that investigated the acoustic characteristics of the sites and offer valuable insights into the auditory experiences at rock art sites, shedding light on their unique acoustic properties.

## Introduction

Rock art sites are places of archaeological interest where human-made markings can be found on natural surfaces. An aggregation site is a particular type of site where hunter-gatherers and agricultural and pastoral groups gathered periodically to conduct many types of transactions in a context where social and ritual activities were of key importance, often involving production of rock art (Bahn, 1982; Moure-Romanillo, 1994). One of their characteristics is that they are surrounded by other sites denominated as “satellites”, characterized by a smaller size and a lesser amount of archaeological remains and rock art motifs. The *Cuevas de la Araña* rock art site, located in Bicorp (Spain), has been identified as one of such aggregation sites.

In July 2021, a fieldwork campaign was conducted, during which acoustic measurements were performed on the site of *Cuevas de la Araña*, its satellites (with a lesser amount and diversity of rock art), and some mountain shelters without rock art (for further information about the sites included in the study and the methodology used, please refer to Santos da Rosa et al., 2023). In this study, a widely recognized technique to capture the acoustic characteristics of an environment was employed, consisting of the recording of impulse responses (Farina, 2007). The objective of this study was to answer the following research question: do aggregation sites have acoustics that would potentiate the perceptual impact of social and ritual activities carried out in these spaces? This seems plausible since rock art in aggregation sites is produced in ritualistic contexts involving singing and dancing (Díaz-Andreu et al., 2021; Domingo et al., 2020). The acoustic analysis showed that the acoustic peculiarities of *Cuevas de la Araña* regarding reverberation could have intensified the sensory effect and emotional impact of the ceremonies likely performed with musical accompaniment (Santos da Rosa et al., 2023).

The study of acoustics by recording impulse responses not only facilitates the examination of acoustic properties from a physical standpoint, but also enables creating auralizations (Kleiner & Dalenbäck, 1993), akin to visualizations, which provide the means to recreate an acoustic environment in a laboratory setting. This allows us to explore rock art sites through the lens of auditory perception. The present study aims to investigate, via a listening test, the extent to which the acoustics of *Cuevas de la Araña* can interfere in modern-day listeners’ perception of sound, using auralizations generated with the impulse responses collected in *Cuevas de la Araña* and its surrounding shelters. Additionally, in accordance with prior investigations in the field of psychoacoustics related to rock art sites (López-Mochales, Aparicio-Terrés, et al., 2023), this study aims to explore variations in the auditory perception of sites featuring rock art in comparison to sites devoid of such art.

## Methods

The listening test consisted of a comparison between pairs of auralizations of different shelters, including the main site of *Cuevas de la Araña* and other surrounding sites. Ten healthy human volunteers –seven males and three females of ages between 25 and 34 years old-took part on the listening test. The exclusion criteria for participants included hearing impairments, psychiatric or neurological illnesses, ages below 18 or above 35 years and consumption of drugs or pharmaceuticals acting on the central nervous system. The test was conducted in November 2022 in the multi-channel chamber of the Aalto Acoustics Lab (Aalto University, Finland). This facility consists of an anechoic chamber equipped with an array of 45 loudspeakers surrounding the listener’s position.

Six impulse responses (IRs) were studied in this test: three measured in sites with rock art (from now on, referred to as *art+*) and three recorded in nearby mountain shelters of similar geomorphology without any traces of rock art (from now on, referred to as *art-*). The *art+* IRs included one recorded in the main shelter of *Cuevas de la Araña*, and two recorded in two satellite sites located in the same area: *Abrigo del Voro* and *Balsa de Calicanto*. All the IRs were collected during the fieldwork campaign of the Artsoundscapes project in Valencia (Spain) during July 2021, taking the requirements included in the ISO 3382-1 (*Acoustics. Measurement of Room Acoustic Parameters. Part 1: Performance Spaces*, 2009), adapting to the particularities of the mountain shelters (Alvarez-Morales et al., 2023; López-Mochales, Alvarez-Morales, et al., 2023; Till, 2020).

In order to measure the IRs, a dodecahedral loudspeaker (IAG DD4 mini dodecahedral loudspeaker) and a 3^rd^ order Ambisonics spherical microphone array (Zylia ZM-1) were used to reproduce and record the excitation signal: a 12-seconds long exponential sine-sweep starting from 50Hz to 20kHz. The recordings of the Ambisonics microphone were post-processed with the SIMO script (Benitez-Aragon et al., 2023) programmed in Matlab R2022a to obtain the spatial IRs. Additionally, the monaural IRs were registered using an omnidirectional microphone (micW n201) and the software tool EASERA 1.2 (*EASERA: Electronic and Acoustic System Evaluation and Response Analysis*, n.d.) to extract the monaural acoustic parameters that characterize the spaces studied (see Table 1).

**Table 1.** Acoustic parameters (averaged from 250Hz to 4kHz) calculated from the monaural IRs selected for the present study: *Cuevas de la Araña* (CA), *Balsa de Calicanto* (BC), *Abrigo del Voro* (VO) and the three *art-* shelters, or shelters without rock art (N1, N2 and N3).

| | $T^*_{20m}$ (s) | $C_{80m}$ (dB) | $D_{50m}$ | $T_{Sm}$ |
| --- | --- | --- | --- | --- |
| CA | 0.85 | 15.70 | 0.96 | 15.67 |
| BC | 0.31 | 18.78 | 0.96 | 17.33 |
| VO | 0.23 | 23.90 | 0.98 | 13.04 |
| N1 | 0.23 | 24.58 | 0.99 | 9.64 |
| N2 | 0.13 | 31.30 | 0.99 | 9.54 |
| N3 | 0.15 | 29.52 | 0.99 | 10.43 |

\*Values of reverberation parameters might not be precise in all the sites due to non-linearities in their decay curves, and therefore should be interpreted with caution.

Multiple source-receiver combinations were considered at each shelter to collect a representative set of impulse responses (IRs) for each site; of these, one from each site was selected to conduct the listening test. Figure 1 illustrates the locations of the source and receiver used for recording of the IRs employed in the test. To the extent feasible, the sound source was positioned at a height of 1.50m, and the receiver was set at a height ranging between 1.40m to 1.50m. These heights were chosen to closely approximate the average ear level of a standing speaker or listener, taking into account the spatial constraints encountered at most of the sites. Both the source and receiver were positioned with a minimum separation of 1m from any surface and at least 5m apart from each other. In the *art+* locations, the source was situated in front of the paintings. When the available space allowed, the receiver was positioned on the platform facing the wall, considering the expected position of a hypothetical audience during activities related to the creation of rock art.

**Figure 1.**
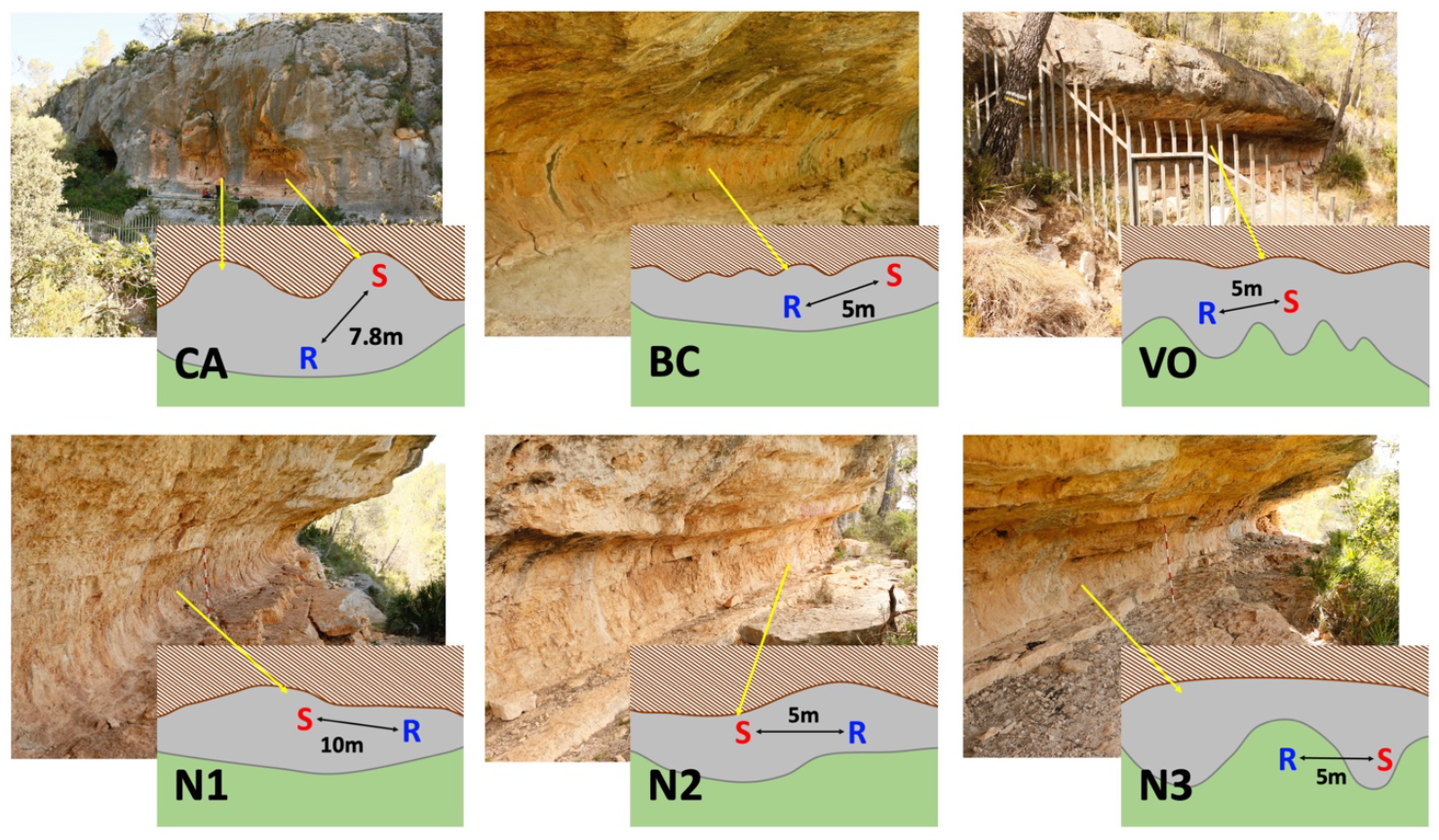
Pictures and schematic representations of the mountain shelters: *Cuevas de la Araña* (CA), *Balsa de Calicanto* (BC), *Abrigo del Voro* (VO) and the three *art-* shelters, or shelters without rock art (N1, N2 and N3). Yellow arrows indicate the correspondence of particular spots of the shelter in the schematic drawings. Letters indicate the position of the source (S) and receiver (R) devices in the recordings of each impulse response, with respect from the mountain walls (brown dashed area) and the platform (grey area). Numbers indicate the distance between the source and receiver in each case.

The stimulus set assembled for this listening test comprised five sound excerpts of 20 seconds duration, convolved with the six impulse responses described above. The sounds were an excerpt of a female singer humming a popular Catalan song entitled *La dama d’Aragó*, a male singer humming the same popular song, the same female singer humming an excerpt of the song *Payphone* by the band Maroon 5, the male singer humming the same excerpt of *Payphone*, and a reconstruction of a shamanic Saami drum (Zachrisson, 1991) beating at 120 bpm. All the sound excerpts were recorded in anechoic conditions and in uncompressed mono format (available as Supp. Mat.). These were retrieved from the ERC Artsoundscapes project’s own repository of anechoic sounds (López-Mochales & Jiménez-Pasalodos, 2021). Each sound excerpt was convolved with each one of the six impulse responses, resulting in six different versions of each sound excerpt, and a total of 30 different auralizations. The convolutions were performed using the MatrixConv plugin from the SPARTA suite (McCormack & Politis, 2019). The signals were decoded for the array of 45 loudspeakers of the multi-channel chamber of the Aalto Acoustics Lab, using the AmbiDEC plugin from the SPARTA suite (McCormack & Politis, 2019). The auralizations were rendered to ensure that the sound source consistently remained in front of the listener. The volume of the sounds was equalized by quantifying the sound pressure level (SPL) at the central point of the room, where the participant’s head was positioned, using an SPL-meter. The signal levels were then manually aligned to prevent the listeners from fixating solely on noticeable variations in sound intensity, rather than paying attention to other acoustic cues.

The test consisted of 45 trials. On each trial, two auralizations were presented, consisting of the same sound excerpt convolved with one of the *art+* IRs and with one of the *art-* IRs. Using an electronic tablet, participants were allowed to reproduce the excerpts in loop, switching between the two versions, blindly labelled “A” and “B”, as many times as needed (see Fig. 2). They could also press the button “Stop” to stop the sounds. Participants were asked to write, on a separate sheet of paper, at least one and up to four differences perceived between the two versions of the sound excerpt, using words or short expressions of their choice. Once they had written all the differences, they could press the button “Next” on the user interface to move to the following trial. Each one of the ten excerpts was presented convolved with all the possible combinations of *art+* and *art-* IRs. The order of trials was randomized, and the assignation of the *art+* and *art-* auralizations to the A and B labels within trials was also randomized and counterbalanced.

**Figure 2.**
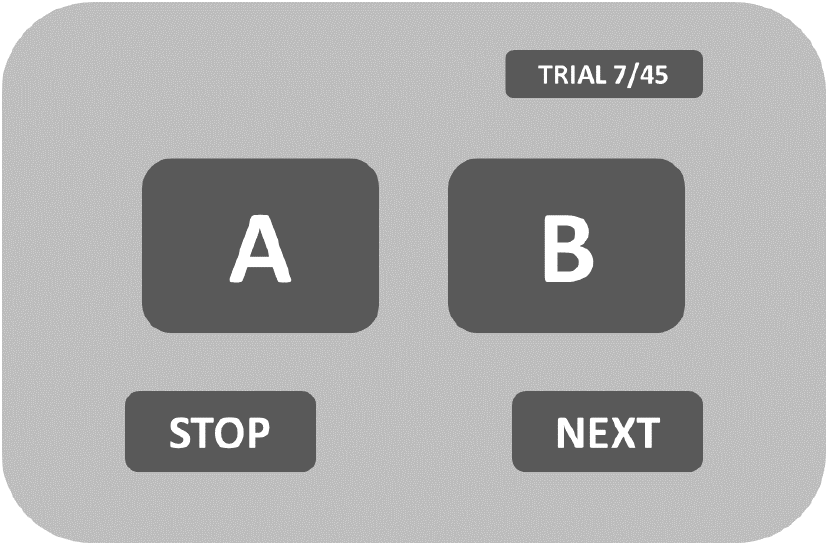
User interface displayed in the electronic tablet.

## Results

A list of 1852 words and short expressions was retrieved from participants’ responses, expressing the differences perceived between the auralizations of the *art+* and *art-* impulse responses.

The classification of the words and expressions was conducted using the R package *tm*, a framework for text mining applications (Feinerer et al., 2008), and consisted of pre-processing, vectorizing and clustering the text. Punctuations and stop words were removed from the list, and it was all converted to lowercase. Then, a document-term matrix was built, to create a numerical representation of the words and expressions (Nguyen, 2013). Finally, the list entries were classified into clusters using the K-Means clustering algorithm (Sterling et al., 2018). The clusters were based on co-occurrence patterns observed in the document-term matrix; words using similar patterns were grouped together. Ten clusters were obtained.

Clusters one to seven included words and phrases associated with the perception of *room size*, the room’s *width*, the prominence of *low frequencies*, the room’s *narrowness*, the *directness* of sound, and the level of *reverberation*, respectively. Clusters eight and nine were excluded from consideration due to their limited size, containing only eight and nine words or expressions, respectively. Cluster number ten contained an assortment of terms that did not align with any other defined category. Given the similarity in meaning, clusters two and four, both pertaining to perceived spatial width, were grouped as a single semantic category in subsequent analyses. Additionally, although cluster number six exclusively featured the word *far* and its variations, the words *close* and *closer* were prevalent in cluster number ten, occurring 182 times. For further analysis, these terms were integrated into the same semantic category as the words from cluster number six. Words and phrases within each semantic category were categorized into opposing ends, positive and negative, of a bipolar scale. As a result, words in the semantic category of *room size* were categorized into those indicating the sound was perceived in a larger space and those indicating it was perceived in a smaller space. Table 2 provides an overview of the classification of words and expressions into these semantic categories.

**Table 2.** Classification of words and expressions into semantic categories.

| Semantic categories | Original clusters | Opposite endpoints of bipolar scales |  |
| --- | --- | --- | --- |
|  |  | Negative | Positive |
| *Room size* | Cluster 1 | *Small / smaller* | *Big / bigger / large* |
| *Room width* | Cluster 2, Cluster 4 | *Narrow / narrower* | *Wide / wider* |
| *Low frequencies* | Cluster 3 | *Less lows* | *More lows / lower* |
| *Directness* | Cluster 5 | *Less direct* | *More direct* |
| *Distance* | Cluster 6 and words "close" / "closer" in Cluster 10 | *Close / Closer* | *Farther / farther away* |
| *Reverberation* | Cluster 7 | *Less reverb / less reverberant* | *More reverb / more reverberant* |

A contingency table was created for each individual semantic category, containing the counts of words of each opposite endpoint, assigned by participants to sounds processed through each of the six impulse responses. Subsequently, a Chi-squared test was conducted for each of these contingency tables, to ascertain the presence of a statistically significant relationship between the participants’ usage of positive and negative words to characterize the sounds and the specific impulse responses with which the sounds were convolved. After adjusting the p-values using the Bonferroni method to mitigate type-I errors (Shaffer, 1995), every outcome yielded a significant result (Fig. 3). An effect of the impulse response was observed on the count of positive and negative words assigned to sounds within the semantic categories of *room size* [X^2^(5,58) = 30.818, p < 0.001], *room width* [X^2^(5,76) = 54.264, p < 0.001], *low frequencies* [X^2^(5,85) = 27.316, p < 0.001], *directness* [X^2^(5,20) = 20.000, p < 0.01], *distance* [X^2^(5,211) = 179.820, p < 0.001], and *reverberation* [X^2^(5,393) = 343.680, p < 0.001].

**Figure 3.**
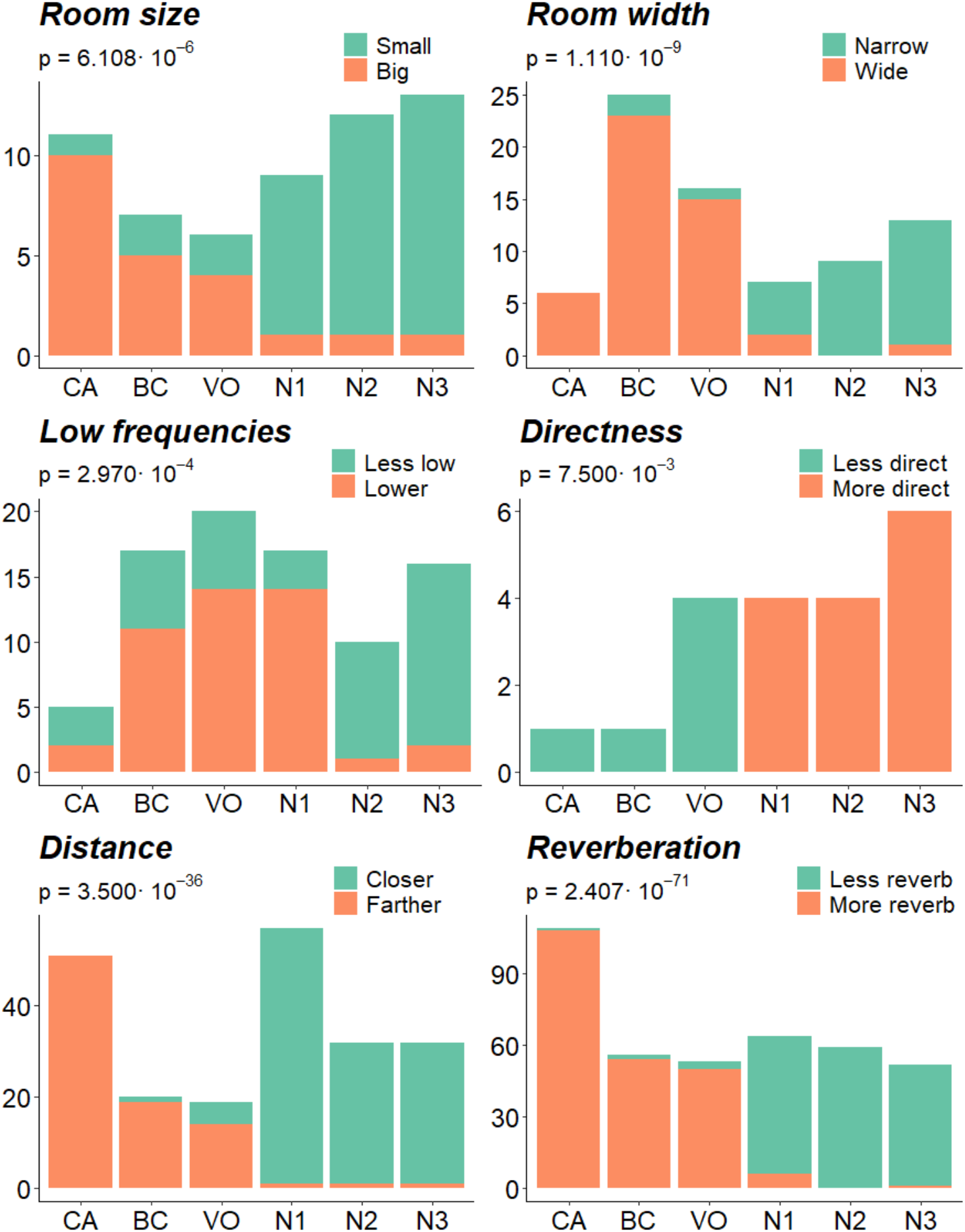
Count of words on the positive (orange) and negative (green) endpoints of each semantic category, assigned by participants to the auralizations of each impulse response: *Cuevas de la Araña* (CA), *Balsa de Calicanto* (BC), *Abrigo del Voro* (VO) and the three shelters without rock art (N1, N2 and N3).

Pearson residuals were computed (Sharpe, 2015) to investigate variations among the outcomes of the different impulse responses, with especial interest in the results of the site of *Cuevas de la Araña*, and to uncover potential patterns that might group the *art+* and *art-* ones (Table 3).

**Table 3.** Pearson residuals of the Chi-square tests performed on the contingency tables containing the count of words in the opposite endpoints of each semantic category, attributed to the auralizations of each impulse response: *Cuevas de la Araña* (CA), *Balsa de Calicanto* (BC), *Abrigo del Voro* (VO) and the three shelters without rock art (N1, N2 and N3).

|  | *Small* | *Big* |  | *Narrow* | *Wide* |
| --- | --- | --- | --- | --- | --- |
| **CA** | -2.2303 | 2.8530 | **CA** | -1.5131 | 1.1886 |
| **BC** | -1.1249 | 1.4390 | **BC** | -2.4411 | 1.9175 |
| **VO** | -0.8934 | 1.1429 | **VO** | -2.0662 | 1.6230 |
| **N1** | 1.0213 | -1.3064 | **N1** | 1.4250 | -1.1194 |
| **N2** | 1.3014 | -1.6648 | **N2** | 3.0034 | -2.3592 |
| **N3** | 1.3839 | -1.7703 | **N3** | 3.1606 | -2.4827 |
|  | *Less low* | *Lower* |  | *Less direct* | *More direct* |
| **CA** | 0.3788 | -0.3656 | **CA** | 1.2780 | -0.8367 |
| **BC** | -0.7683 | 0.7416 | **BC** | 1.2780 | -0.8367 |
| **VO** | -1.1742 | 1.1335 | **VO** | 2.5560 | -1.6733 |
| **N1** | -1.8159 | 1.7529 | **N1** | -1.0954 | 0.7171 |
| **N2** | 1.9016 | -1.8357 | **N2** | -1.0954 | 0.7171 |
| **N3** | 2.2614 | -2.1830 | **N3** | -1.3416 | 0.8783 |
|  | *Closer* | *Farther* |  | *Less reverb* | *More reverb* |
| **CA** | -5.4746 | 6.5359 | **CA** | -6.8030 | 6.0639 |
| **BC** | -3.1367 | 3.7447 | **BC** | -4.5777 | 4.0804 |
| **VO** | -1.8452 | 2.2029 | **VO** | -4.2248 | 3.7658 |
| **N1** | 3.8880 | -4.6417 | **N1** | 5.5727 | -4.9672 |
| **N2** | 2.8120 | -3.3571 | **N2** | 6.4328 | -5.7339 |
| **N3** | 2.8120 | -3.3571 | **N3** | 5.8307 | -5.1973 |

The semantic category of *low frequencies* is the only one in which we did not observe a consistency in the sign of the residuals within each group of impulse responses, *art+* and *art-*. Additionally, the absolute residuals for the word counts associated with the *Cuevas de la Araña* site are notably small.

In contrast, for all the remaining semantic categories, a uniform pattern was observed in the sign of the residuals within each group of impulse responses, *art+* and *art-*. Participants described the *art+* auralizations –including those created with the IRs gathered from the main site of *Cuevas de la Araña* and the satellite sites of *Balsa de Calicanto* and *Abrigo del Voro*-as *bigger, wider, less direct, farther* and *more reverberant* than the *art-* ones.

Examining the absolute magnitude of the residuals serves as an indicator of the disparity between observed and expected counts within a particular cell of the contingency table (Sharpe, 2015). In the tables associated with the semantic categories of *size, distance* and *reverberation* the largest absolute magnitudes of the residuals were observed in the cells corresponding to the IR of *Cuevas de la Araña*.

## Discussion

In the present study, a listening test was conducted to examine the differences in the acoustic perception of a series of auralizations generated using impulse responses collected from natural mountain shelters located in Bicorp, within the Comunitat Valenciana region of Spain. The primary objectives were to determine whether the acoustic qualities of the *Cuevas de la Araña* rock art site were distinctly perceived compared to other nearby sites in the same vicinity. Additionally, the study aimed to identify the specific dimensions of perception where listeners discerned disparities. As a secondary inquiry, the research sought to ascertain whether differences in perception existed between sites with rock art paintings, namely, *Cuevas de la Araña* and two satellite sites, *Balsa de Calicanto* and *Abrigo del Voro*, and those that did not present any traces of rock art, despite sharing similar geomorphological features. The listening test involved a comparison of paired stimuli, consistently contrasting an auralization from a site with rock art and one from a site without rock art. The findings revealed perceptual differences between sites with and without rock art in the dimensions of apparent size of the space, width, sound directness, source-to-listener distance, and reverberation. Notably, the most pronounced differences in perception were observed in the dimensions of apparent size, distance from the source, and reverberation when comparing auralizations from *Cuevas de la Araña* with those from non-painted sites.

The inconsistent pattern within the semantic category of low frequencies implies that this particular acoustic feature may not have been a key determinant when assessing the appropriateness of a shelter for rock art production and related activities. In the context of the other perceptual dimensions that were examined, significant differences were found between sites featuring rock art and those lacking it. As indicated by the outcomes of the acoustic analysis (Santos da Rosa et al., 2023), the impulse responses gathered at *Cuevas de la Araña* exhibit some reflections from the canyon where it is located that lead to the certain increase in reverberation in comparison to those collected in the satellite sites and the control sites lacking rock art. Nevertheless, participants characterized the auralizations derived from the three rock art sites -*Cuevas de la Araña* and the two satellite sites, *Balsa de Calicanto* and *Abrigo del Voro*-as having a heightened sense of reverberation when compared to the controls. This enhanced sense of reverberation played a pivotal role in influencing the perception of spatial characteristics, imparting a notion of increased room size and width (Cabrera et al., 2005). In comparison to other environments perceived as acoustically drier, the sense of reverberation created the perceptual illusion of an augmented distance between the sound source and the listener, a phenomenon previously noted (Bronkhorst & Houtgast, 1999; Kolarik et al., 2016). Ultimately, within such reverberant environments, by definition, sounds are inherently perceived as less direct, due to the blending of direct sound with surface reflections. Because of the reverberation of the impulse response from *Cuevas de la Araña* being the longest of those studied, the differences in the perceptual dimensions of room size, distance and reverberation become notably pronounced when contrasting the auralizations of this location with those of unpainted sites.

In the present study, a sensory descriptive analysis methodology was employed (Stone et al., 2012), wherein participants were asked to indicate the differences perceived between pairs of stimuli, employing their individual vocabulary. Subsequently, the ascribed descriptors were classified through a hierarchical clustering method. To avoid the manual scrutiny and categorization of text, Natural Language Processing techniques are typically used for the analysis of open-ended questionnaire data (Hamzah & Widyastuti, 2016; Inui et al., 2003; Ramachandran, 2018; Spasić et al., 2019). Nevertheless, these techniques are primarily tailored for the analysis of extensive text corpora, whereas our study involved a short list of singular words and short expressions. Consequently, we adopted a hierarchical classification approach akin to that adopted by Garcia-Constantino et al. (2012). Nonetheless, Garcia-Constantino et al. employed a top-down methodology, using a labeled document training set. In contrast, in our study, the words were classified in a bottom-up fashion, relying on orthographic similarity. This approach was chosen to eliminate human judgment in the labeling of categories. However, future studies may benefit from other sophisticated methods for word classification, considering criteria beyond orthography. This could enhance efficiency by accounting for synonyms and semantically related words. The emergence, in our analysis, of a large cluster of words that did not fit into any other category likely results from the employed classification methodology.

Despite the significant results obtained, it is imperative to acknowledge certain limitations that warrant consideration when interpreting our findings. A primary concern lies in the inherent reductionism associated with our research, which focuses on rock art sites and its associated activities conducted millennia ago, examined through the lens of modern resources and knowledge. Concerning this issue, the selection of sounds for our listening test was based on our current understanding, including female and male singing as representatives of known activities (Díaz-Andreu et al., 2021; Domingo et al., 2020), and drumming as an impulse-like sound that would put into manifest certain characteristics such as spatial impression (Pulkki et al., 2004). This leaves open the possibility that other sounds were historically produced, potentially introducing unexplored acoustic cues. Moreover, our auralizations were static and failed to account for potential variations in the position of sound sources and receivers during the activities taken place in prehistoric times, thereby overlooking any acoustic effects that may have arisen from such movements (Ackermann et al., 2019). In the context of the limitations arising from the method of IRs gathering and convolution, the directivity of each sound source –thus, the singers and the drum-was also neglected. Lastly, the exclusive isolation of the aural component in our listening test raises questions about the potential synergy with other sensory inputs, such as the visual, which could have influenced the perception of certain acoustic cues (Udesen et al., 2015). These limitations underscore the need for caution when extrapolating our results to the broader context of rock art sites and their acoustic properties.

## Conclusions

The results of our study provide valuable insights into the perceptual differences between auralizations produced with the impulse responses from rock art sites and sites without traces of rock art. Our findings revealed a compelling association between the impulse responses and the usage of positive and negative words within the semantic categories of room size, width, presence of low frequencies, directness of sound, distance from the sound source and reverberation. Notably, the semantic category of low frequencies exhibited an inconsistent pattern. However, the other results consistently portrayed the auralizations from the rock art sites as larger, wider, less direct, farther, and more reverberant compared to the controls. Examination of the absolute residuals underscored the substantial influence of the *Cuevas de la Araña* impulse response in the categories of size, distance, and reverberation.

These findings suggest that acoustics likely played a significant role in the selection of suitable locations for rock art creation and associated activities. Furthermore, the acoustic properties of *Cuevas de la Araña* may have contributed to its repeated visits and activities settings that eventually led to its recognition as an aggregation site. The results, which confirm participants’ discernment of substantial differences in auralizations between this site and others, align with the previous acoustic analysis by Santos da Rosa et al. (2023), which concluded that the acoustic characteristics of *Cuevas de la Araña* could enhance the appreciation of activities accompanied by music.

## Acknowledgements

We extend our sincere gratitude to everyone involved in managing and conducting the fieldwork campaign, and everyone responsible for registering and analyzing the impulse responses. Gratitude also to the Aalto Acoustics Lab at Aalto University, Finland, for their vital support in implementing our experiment in their laboratory.

## Funding

This work is part of the ERC Artsoundscapes project (grant agreement no. 787842) that has received funding from the European Research Council (ERC) under the European Union’s Horizon 2020 research and innovation program. PI: MD-A. CE was also supported by the Generalitat de Catalunya SGR2017-974, de María de Maeztu Center of Excellence (Institute of Neurosciences, University of Barcelona) CEX2021-001159-M, Ministry of Science and Innovation, and the ICREA Acadèmia Distinguished Professorship Award.

## Notes

https://zenodo.org/records/11046468

### Competing Interest Statement

The authors have declared no competing interest.

